# Comprehensive Analysis of *HEXB* Protein Reveal Forty Two Novel nsSNPs That May Lead to Sandhoff disease (SD) Using Bioinformatics

**DOI:** 10.1101/853077

**Authors:** Tebyan A. Abdelhameed, Mosab M. Gasmelseed, Mujahed I. Mustafa, Dina N. Abdelrahman, Fatima A. Abdelrhman, Mohamed A. Hassan

**Author notes:** Corresponding author: Tebyan A. Abdelhameed.

## Abstract

**Background:** Single Nucleotide Polymorphisms (SNPs) in the *HEXB* gene are associated with a neurodegenerative disorder called Sandhoff disease (SD) (GM2 gangliosidosis-O variant). This study aimed to predict the possible pathogenic SNPs of this gene and their impact on the protein using different bioinformatics tools.

**Methods:** SNPs retrieved from the NCBI database were analyzed using several bioinformatics tools. The different algorithms collectively predicted the effect of single nucleotide substitution on both structure and function of beta subunit beta subunit of both hexosaminidase A and hexosaminidase B proteins.

**Results:** Forty nine mutations were found to be extremely damaging to the structure and function of the HEXB gene protein.

**Conclusion:** According to this study, forty two novel nsSNP in *HEXB* are predicted to have possible role in Sandhoff disease using different bioinformatics tools, beside two SNPs found to have effect on miRNAs binding site affecting expression of *HEXB* gene. Our findings may assist in genetic study and diagnosis of Sandhoff disease.

## 1. INTRODUCTION

Sandhoff disease (SD) also called (GM2 gangliosidosis-O variant) is a fatal rare autosomal recessive lysosomal storage disease of sphingolipid (GM2 ganglioside GM2) metabolism resulting from the deficiency of β-hexosaminidase (HexB) (1-5). The estimated carrier frequency of SD with a high incidence among certain isolated communities and ethnic groups around the world such as in Saskatchewan at rate 1:15 (1, 6). It is caused by deficiency of N-acetyl-β-hexosaminidase (Hex) resulting in pathological accumulation of GM2 ganglioside in lysosomes of the central nervous system (CNS) and progressive neurodegeneration. Currently, there is no treatment for SD. (2, 7) Affected individuals present with a wide spectrum of clinical manifestations, ranging from psychomotor impairment and death in the infantile form to motor neuron disease and autonomic dysfunction in the adult form(8) The disorder is classified according to the age of onset, as infantile, juvenile and adult form.(5) The most common and severe form of Sandhoff disease becomes apparent in infancy,(9-13) The clinical features of juvenile SD include ataxia, dysarthria and cerebellar atrophy, develop early and severe sensory loss in addition to chronic motor neuron disease and cerebellar ataxia(14) While the clinical features of adult SD include progressive muscle cramps, as well as wasting and weakness of the legs with onset after age 20. They also can show intention tremor of the upper extremities and dysarthria.(8)

The *HEXB* gene provides instructions for making a protein that is a subunit of two related enzymes; beta-hexosaminidase A and beta-hexosaminidase B. play a critical role in the central nervous system, These enzymes are found in lysosomes. Within lysosomes, the enzymes break down sphingolipids, oligosaccharides, and molecules that are linked to sugars such as glycoproteins (15). *HEXB* gene is localized on 5q13.3, which is the long (q) arm of chromosome 5 at position 13.3. The disease is caused by mutation in *HEXB* encoding the β-subunit of β-hexosaminidase A. β-Hexosaminidase A exists as a heterodimer consisting of α- and β-subunits (16). Numerous studies in the past have shown that SNPs are responsible of mutations that lead to cause Sandhoff disease (16, 17). Single nucleotide polymorphisms (SNPs) are an important source of human genome variability. Non-synonymous SNPs occurring in coding regions result in single amino acid polymorphisms (SAPs) that may affect protein function and lead to pathology (18). Functional variations can have deleterious or neutral effects on protein structure or function (19) Damaging effects might include destabilization of protein structure, altering gene regulation (20) affecting protein charge, geometry, hydrophobicity(21), stability, dynamics, translation and inter/intra protein interactions. (22, 23), hence structural integrity of cells comes under risk (24). Thus it can be avowed that nsSNPs might get linked with many human diseases because of these missense SNPs.(25) The aim of this study was to identify the possible pathogenic SNPs in *HEXB* gene using in silico prediction software, and to determine the structure, function and regulation of their respective proteins. This is the first study which covers an extensive in silico analysis of nsSNPs of HEXB protein hence this work might be useful in future in developing precision medicines for the treatment of diseases caused by these genomic variations. (26)

## 2. MATERIALS AND MEYHODS

### 2.1 Data mining

The data on human *HEXB* gene was collected from National Center for Biological Information (NCBI) web site (27). The SNP information (protein accession number and SNP ID) of the MEFV gene was retrieved from the NCBI dbSNP (http://www.ncbi.nlm.nih.gov/snp/) and the protein sequence was collected from Swiss Prot databases (http://expasy.org/). (28)

### 2.2 SIFT

SIFT is a sequence homology-based tool (29) that sorts intolerant from tolerant amino acid substitutions and predicts whether an aminoacid substitution in a protein will have a phenotypic Effect. Considers the position at which the change occurred and the type of amino acid change. Given a protein sequence, SIFT chooses related proteins and obtains an alignment of these proteins with the query. Based on the amino acids appearing at each position in the alignment, SIFT calculates the probability that an amino acid at a position is tolerated conditional on the most frequent amino acid being tolerated. If this normalized value is less than a cutoff, the substitution is predicted to be deleterious. SIFT scores <0.05 are predicted by the algorithm to be intolerant or deleterious amino acid substitutions, whereas scores >0.05 are considered tolerant. It is available at (http://sift.bii.a-star.edu.sg/).

### 2.3 Polyphen-2

It is a software tool (30) to predict possible impact of an amino acid substitution on both structure and function of a human protein by analysis of multiple sequence alignment and protein 3D structure, in addition it calculates position-specific independent count scores (PSIC) for each of two variants, and then calculates the PSIC scores difference between two variants. The higher a PSIC score difference, the higher the functional impact a particular amino acid substitution is likely to have. Prediction outcomes could be classified as probably damaging, possibly damaging or benign according to the value of PSIC as it ranges from (0_1); values closer to zero considered benign while values closer to 1 considered probably damaging and also it can be indicated by a vertical black marker inside a color gradient bar, where green is benign and red is damaging. nsSNPs that predicted to be intolerant by Sift has been submitted to Polyphen as protein sequence in FASTA format that obtained from UniproktB /Expasy after submitting the relevant ensemble protein (ESNP) there, and then we entered position of mutation, native amino acid and the new substituent for both structural and functional predictions. PolyPhen version 2.2.2 is available at (http://genetics.bwh.harvard.edu/pph2/index.shtml).

### 2.4 Provean

Provean is a software tool (31) which predicts whether an amino acid substitution or indel has an impact on the biological function of a protein. it is useful for filtering sequence variants to identify nonsynonymous or indel variants that are predicted to be functionally important. It is available at (https://rostlab.org/services/snap2web/).

### 2.5 SNAP2

Functional effects of mutations are predicted with SNAP2 (32). SNAP2 is a trained classifier that is based on a machine learning device called “neural network”. It distinguishes between effect and neutral variants/non-synonymous SNPs by taking a variety of sequence and variant features into account. The most important input signal for the prediction is the evolutionary information taken from an automatically generated multiple sequence alignment. Also structural features such as predicted secondary structure and solvent accessibility are considered. If available also annotation (i.e. known functional residues, pattern, regions) of the sequence or close homologs are pulled in. In a cross-validation over 100,000 experimentally annotated variants, SNAP2 reached sustained two-state accuracy (effect/neutral) of 82% (at an AUC of 0.9). In our hands this constitutes an important and significant improvement over other methods. It is available at (https://rostlab.org/services/snap2web/).

### 2.6 PHD-SNP

An online Support Vector Machine (SVM) based classifier, is optimized to predict if a given single point protein mutation can be classified as disease-related or as a neutral polymorphism, it is available at: (http://snps.biofold.org/phd-snp/phdsnp.html).

### 2.7 SNP& Go

SNPs&GO is an accurate method that, starting from a protein sequence, can predict whether a variation is disease related or not by exploiting the corresponding protein functional annotation. SNPs&GO collects in unique framework information derived from protein sequence, evolutionary information, and function as encoded in the Gene Ontology terms, and outperforms other available predictive methods. (33) It is available at (http://snps.biofold.org/snps-and-go/snps-and-go.html)

### 2.8 P-Mut

PMUT a web-based tool (34) for the annotation of pathological variants on proteins, allows the fast and accurate prediction (approximately 80% success rate in humans) of the pathological character of single point amino acidic mutations based on the use of neural networks. It is available at (http://mmb.irbbarcelona.org/PMut).

### 2.9 I-Mutant 3.0

I-Mutant 3.0 Is a neural network based tool (35) for the routine analysis of protein stability and alterations by taking into account the single-site mutations. The FASTA sequence of protein retrieved from UniProt is used as an input to predict the mutational effect on protein stability. It is available at (http://gpcr2.biocomp.unibo.it/cgi/predictors/I-Mutant3.0/I-Mutant3.0.cgi).

### 2.10 Project Hope

Online software is available at: (http://www.cmbi.ru.nl/hope/method/). It is a web service where the user can submit a sequence and mutation. The software collects structural information from a series of sources, including calculations on the 3D protein structure, sequence annotations in UniProt and prediction from other software. It combines this information to give analysis for the effect of a certain mutation on the protein structure. HOPE will show the effect of that mutation in such a way that even those without a bioinformatics background can understand it. It allows the user to submit a protein sequence (can be FASTA or not) or an accession code of the protein of interest. In the next step, the user can indicate the mutated residue with a simple mouse click. In the final step, the user can simply click on one of the other 19 amino acid types that will become the mutant residue, and then full report well is generated (36).

### 2.12 Raptor X

RaptorX (http://raptorx.uchicago.edu/): It is a web server predicting structure property of a protein sequence without using any templates. It outperforms other servers, especially for proteins without close homologs in PDB or with very sparse sequence profile. The server predicts tertiary structure (37)

### 2.11 UCSF Chimera (University of California at San Francisco)

UCSF Chimera (https://www.cgl.ucsf.edu/chimera/) is a highly extensible program for interactive visualization and analysis of molecular structures and related data, including density maps, supramolecular assemblies, sequence alignments, docking results, trajectories, and conformational ensembles. High-quality images and animations can be generated. Chimera includes complete documentation and several tutorials. Chimera is developed by the Resource for Biocomputing, Visualization, and Informatics (RBVI), supported by the National Institutes of Health (P41-GM103311).(38)

### 2.12 PolymiRTS

PolymiRTS is a software used to predict 3UTR (un-translated region) polymorphism in microRNAs and their target sites available at (http://compbio.uthsc.edu/miRSNP/). It is a database of naturally occurring DNA variations in mocriRNAs (miRNA) seed region and miRNA target sites. MicroRNAs pair to the transcript of protein coding genes and cause translational repression or mRNA destabilization. SNPs in microRNA and their target sites may affect miRNA-mRNA interaction, causing an effect on miRNA-mediated gene repression, PolymiRTS database was created by scanning 3UTRs of mRNAs in human and mouse for SNPs in miRNA target sites. Then, the effect of polymorphism on gene expression and phenotypes are identified and then linked in the database. The PolymiRTS data base also includes polymorphism in target sites that have been supported by a variety of experimental methods and polymorphism in miRNA seed regions. (39)

### 2.13 GeneMANIA

It is gene interaction software that finds other genes which is related to a set of input genes using a very large set of functional association data. Association data include protein and genetic interactions, pathways, co-expression, co-localization and protein domain similarity. GeneMANIA also used to find new members of a pathway or complex, find additional genes you may have missed in your screen or find new genes with a specific function, such as protein kinases. available at (https://genemania.org/) (40).

## 3. RESULTS

### 3.1 The functional effect of Deleterious and damaging nsSNPs of HEXB by SIFT, Provean PolyPhen-2, and SNAP2

**Table (1):** Damaging or Deleterious or effect nsSNPs associated variations predicted by SIFT, Provean PolyPhen-2, and SNAP2 softwares:

| dbSNP rs# | SUB | SIFT prediction | Score | Polyphen prediction | Score | Prediction (cutoff= -2.5) | PROVEAN score | SNAP2 prediction | Score |
| --- | --- | --- | --- | --- | --- | --- | --- | --- | --- |
| rs766282406 | W166C | D | 0 | probably damaging | 1 | Deleterious | -12.526 | effect | 31 |
| rs1205668888 | G167R | D | 0 | probably damaging | 1 | Deleterious | -7.708 | effect | 95 |
| rs951616685 | G167A | D | 0 | probably damaging | 1 | Deleterious | -5.781 | effect | 84 |
| rs759538325 | R170Q | D | 0 | probably damaging | 1 | Deleterious | -3.654 | effect | 72 |
| rs976468812 | F175C | D | 0 | probably damaging | 1 | Deleterious | -7.708 | effect | 4 |
| rs398123449 | D196A | D | 0 | probably damaging | 1 | Deleterious | -7.708 | effect | 63 |
| rs752621035 | R199G | D | 0 | probably damaging | 1 | Deleterious | -6.745 | effect | 75 |
| rs762821794 | G204E | D | 0 | probably damaging | 1 | Deleterious | -7.708 | effect | 85 |
| rs761538803 | R211T | D | 0 | probably damaging | 1 | Deleterious | -5.779 | effect | 90 |
| rs367963796 | L223P | D | 0 | probably damaging | 1 | Deleterious | -6.643 | effect | 77 |
| rs1050332517 | N232H | D | 0 | probably damaging | 1 | Deleterious | -4.873 | effect | 61 |
| rs1375560334 | L234R | D | 0 | probably damaging | 1 | Deleterious | -4.4 | effect | 65 |
| rs1258632902 | H237R | D | 0 | probably damaging | 1 | Deleterious | -7.797 | effect | 77 |
| rs1301523493 | P245L | D | 0 | probably damaging | 1 | Deleterious | -9.663 | effect | 52 |
| rs771103635 | L254S | D | 0 | probably damaging | 1 | Deleterious | -5.846 | effect | 45 |
| rs373979283 | Y266D | D | 0 | probably damaging | 1 | Deleterious | -9.403 | effect | 92 |
| rs398123450 | Y266C | D | 0 | probably damaging | 1 | Deleterious | -8.495 | effect | 68 |
| rs750645495 | E288Q | D | 0 | probably damaging | 1 | Deleterious | -2.806 | effect | 84 |
| rs866283559 | D290N | D | 0 | probably damaging | 1 | Deleterious | -4.71 | effect | 80 |
| rs1200101608 | G293R | D | 0 | probably damaging | 1 | Deleterious | -7.536 | effect | 87 |
| rs1439544901 | W298C | D | 0 | probably damaging | 1 | Deleterious | -11.813 | effect | 24 |
| rs1189912520 | G301D | D | 0 | probably damaging | 1 | Deleterious | -6.585 | effect | 50 |
| rs1252744810 | L305P | D | 0 | probably damaging | 1 | Deleterious | -6.474 | effect | 74 |
| rs773315856 | F344S | D | 0 | probably damaging | 1 | Deleterious | -7.676 | effect | 70 |
| rs751352339 | H350D | D | 0 | probably damaging | 1 | Deleterious | -8.621 | effect | 92 |
| rs781244479 | D354A | D | 0 | probably damaging | 1 | Deleterious | -7.676 | effect | 85 |
| rs781244479 | D354G | D | 0 | probably damaging | 1 | Deleterious | -6.724 | effect | 91 |
| rs1245724719 | C360Y | D | 0 | probably damaging | 1 | Deleterious | -10.553 | effect | 53 |
| rs779364743 | W361R | D | 0 | probably damaging | 1 | Deleterious | -13.153 | effect | 87 |
| rs753450688 | W405R | D | 0 | probably damaging | 1 | Deleterious | -13.495 | effect | 95 |
| rs984639297 | P468H | D | 0 | probably damaging | 1 | Deleterious | -8.462 | effect | 56 |
| rs760333736 | G483D | D | 0 | probably damaging | 1 | Deleterious | -6.504 | effect | 80 |
| rs760333736 | G483V | D | 0 | probably damaging | 1 | Deleterious | -8.362 | effect | 80 |
| rs1447056657 | W489R | D | 0 | probably damaging | 1 | Deleterious | -13.008 | effect | 95 |
| rs754180181 | E491G | D | 0 | probably damaging | 1 | Deleterious | -6.504 | effect | 86 |
| - | V493G | D | 0 | probably damaging | 1 | Deleterious | -6.504 | effect | 54 |
| rs121907985 | P504S | D | 0 | probably damaging | 1 | Deleterious | -7.294 | effect | 91 |
| - | R505W | D | 0 | probably damaging | 1 | Deleterious | -7.292 | effect | 90 |
| rs1229725676 | E511G | D | 0 | probably damaging | 1 | Deleterious | -6.382 | effect | 73 |
| rs778501777 | L513H | D | 0 | probably damaging | 1 | Deleterious | -6.382 | effect | 53 |
| rs778501777 | L513P | D | 0 | probably damaging | 1 | Deleterious | -6.382 | effect | 72 |
| rs1417317737 | W514C | D | 0 | probably damaging | 1 | Deleterious | -11.853 | effect | 77 |
| rs760366178 | R528I | D | 0 | probably damaging | 1 | Deleterious | -7.294 | effect | 77 |
| rs745454291 | L529R | D | 0 | probably damaging | 1 | Deleterious | -5.371 | effect | 69 |
| rs764552042 | R533C | D | 0 | probably damaging | 1 | Deleterious | -7.294 | effect | 63 |
| rs727503960 | C534F | D | 0 | probably damaging | 1 | Deleterious | -10.029 | effect | 23 |
| rs749646826 | R539C | D | 0 | probably damaging | 1 | Deleterious | -7.174 | effect | 32 |
| rs769208361 | R539H | D | 0 | probably damaging | 1 | Deleterious | -4.484 | effect | 61 |
| rs727503961 | C551Y | D | 0 | probably damaging | 1 | Deleterious | -9.773 | effect | 79 |

\*SUB= Substitution
\*D= Deleterious

### 3.2 Functional analysis of ADAMTS13 gene using Disease Related and pathological effect of nsSNPs by PhD-SNP, SNPs & GO and PMut softwares

**Table (2):** Prediction of Disease Related and pathological effect of nsSNPs by PhD-SNP, SNPs & GO and PMut:

| dbSNP rs# | SUB | SNP&Go Prediction | RI | Probability | PHD-SNP Prediction | RI | Probability | P-mut Prediction | Score |
| --- | --- | --- | --- | --- | --- | --- | --- | --- | --- |
| rs766282406 | W166C | Disease | 3 | 0.645 | Disease | 6 | 0.785 | Disease | 0.78 (88%) |
| rs1205668888 | G167R | Disease | 7 | 0.865 | Disease | 9 | 0.937 | Disease | 0.93 (94%) |
| rs951616685 | G167A | Disease | 6 | 0.805 | Disease | 7 | 0.852 | Disease | 0.80 (89%) |
| rs759538325 | R170Q | Disease | 5 | 0.734 | Disease | 5 | 0.732 | Disease | 0.82 (90%) |
| rs976468812 | F175C | Disease | 6 | 0.779 | Disease | 8 | 0.876 | Disease | 0.93 (94%) |
| rs398123449 | D196A | Disease | 7 | 0.861 | Disease | 8 | 0.9 | Disease | 0.92 (94%) |
| rs752621035 | R199G | Disease | 4 | 0.711 | Disease | 7 | 0.837 | Disease | 0.88 (92%) |
| rs762821794 | G204E | Disease | 7 | 0.846 | Disease | 8 | 0.903 | Disease | 0.85 (91%) |
| rs761538803 | R211T | Disease | 7 | 0.863 | Disease | 9 | 0.932 | Disease | 0.92 (94%) |
| rs367963796 | L223P | Disease | 7 | 0.841 | Disease | 9 | 0.951 | Disease | 0.93 (94%) |
| rs1050332517 | N232H | Disease | 7 | 0.829 | Disease | 9 | 0.94 | Disease | 0.93 (94%) |
| rs1375560334 | L234R | Disease | 9 | 0.934 | Disease | 9 | 0.969 | Disease | 0.92 (94%) |
| rs1258632902 | H237R | Disease | 8 | 0.88 | Disease | 9 | 0.948 | Disease | 0.91 (93%) |
| rs1301523493 | P245L | Disease | 4 | 0.704 | Disease | 6 | 0.795 | Disease | 0.87 (92%) |
| rs771103635 | L254S | Disease | 6 | 0.779 | Disease | 6 | 0.824 | Disease | 0.92 (94%) |
| rs373979283 | Y266D | Disease | 7 | 0.848 | Disease | 9 | 0.946 | Disease | 0.93 (94%) |
| rs398123450 | Y266C | Disease | 7 | 0.835 | Disease | 9 | 0.932 | Disease | 0.93 (94%) |
| rs750645495 | E288Q | Disease | 6 | 0.803 | Disease | 7 | 0.837 | Disease | 0.89 (92%) |
| rs866283559 | D290N | Disease | 4 | 0.711 | Disease | 7 | 0.843 | Disease | 0.82 (90%) |
| rs1200101608 | G293R | Disease | 7 | 0.848 | Disease | 8 | 0.892 | Disease | 0.93 (94%) |
| rs1439544901 | W298C | Disease | 3 | 0.63 | Disease | 5 | 0.773 | Disease | 0.84 (90%) |
| rs1189912520 | G301D | Disease | 3 | 0.659 | Disease | 6 | 0.779 | Disease | 0.76 (88%) |
| rs1252744810 | L305P | Disease | 7 | 0.836 | Disease | 8 | 0.899 | Disease | 0.91 (93%) |
| rs773315856 | F344S | Disease | 6 | 0.797 | Disease | 8 | 0.891 | Disease | 0.78 (88%) |
| rs751352339 | H350D | Disease | 6 | 0.821 | Disease | 8 | 0.898 | Disease | 0.92 (94%) |
| rs781244479 | D354A | Disease | 8 | 0.887 | Disease | 9 | 0.928 | Disease | 0.89 (92%) |
| rs781244479 | D354G | Disease | 7 | 0.857 | Disease | 8 | 0.918 | Disease | 0.93 (94%) |
| rs1245724719 | C360Y | Disease | 2 | 0.596 | Disease | 7 | 0.829 | Disease | 0.68 (85%) |
| rs779364743 | W361R | Disease | 7 | 0.856 | Disease | 9 | 0.936 | Disease | 0.93 (94%) |
| rs753450688 | W405R | Disease | 8 | 0.884 | Disease | 9 | 0.956 | Disease | 0.91 (93%) |
| rs984639297 | P468H | Disease | 4 | 0.703 | Disease | 7 | 0.861 | Disease | 0.77 (88%) |
| rs760333736 | G483D | Disease | 8 | 0.884 | Disease | 9 | 0.949 | Disease | 0.92 (94%) |
| rs760333736 | G483V | Disease | 7 | 0.874 | Disease | 9 | 0.941 | Disease | 0.93 (94%) |
| rs1447056657 | W489R | Disease | 8 | 0.895 | Disease | 9 | 0.97 | Disease | 0.93 (94%) |
| rs754180181 | E491G | Disease | 6 | 0.779 | Disease | 7 | 0.86 | Disease | 0.88 (92%) |
| - | V493G | Disease | 4 | 0.697 | Disease | 6 | 0.822 | Disease | 0.74 (87%) |
| rs121907985 | P504S | Disease | 5 | 0.747 | Disease | 6 | 0.819 | Disease | 0.87 (91%) |
| - | R505W | Disease | 7 | 0.827 | Disease | 8 | 0.893 | Disease | 0.91 (93%) |
| rs1229725676 | E511G | Disease | 4 | 0.68 | Disease | 4 | 0.723 | Disease | 0.87 (91%) |
| rs778501777 | L513H | Disease | 1 | 0.541 | Disease | 3 | 0.644 | Disease | 0.84 (90%) |
| rs778501777 | L513P | Disease | 6 | 0.804 | Disease | 6 | 0.819 | Disease | 0.84 (90%) |
| rs1417317737 | W514C | Disease | 7 | 0.83 | Disease | 8 | 0.891 | Disease | 0.92 (93%) |
| rs760366178 | R528I | Disease | 5 | 0.742 | Disease | 8 | 0.91 | Disease | 0.86 (91%) |
| rs745454291 | L529R | Disease | 4 | 0.696 | Disease | 8 | 0.908 | Disease | 0.85 (91%) |
| rs764552042 | R533C | Disease | 5 | 0.752 | Disease | 7 | 0.873 | Disease | 0.79 (89%) |
| rs727503960 | C534F | Disease | 5 | 0.75 | Disease | 6 | 0.794 | Disease | 0.65 (84%) |
| rs749646826 | R539C | Disease | 4 | 0.707 | Disease | 7 | 0.857 | Disease | 0.86 (91%) |
| rs769208361 | R539H | Disease | 0 | 0.516 | Disease | 4 | 0.69 | Disease | 0.59 (82%) |
| rs727503961 | C551Y | Disease | 3 | 0.672 | Disease | 8 | 0.893 | Disease | 0.88 (92%) |

### 3.3 Prediction of Change in Stability due to Mutation Using I-Mutant 3.0 Server

**Table (3):** Prediction of nsSNPs Impact on Protein structure Stability by I-Mutant

3.3 Prediction of Change in Stability due to Mutation Using I-Mutant 3.0 Server **Table (3):** Prediction of nsSNPs Impact on Protein structure Stability by I-Mutant
| dbSNP rs# | SNP | SVM2 Prediction Effect | DDG Value |
| --- | --- | --- | --- |
| rs766282406 | W166C | Decrease Protein Stability | -1.64 |
| rs1205668888 | G167R | Decrease Protein Stability | -0.9 |
| rs951616685 | G167A | Decrease Protein Stability | -0.68 |
| rs759538325 | R170Q | Decrease Protein Stability | -0.84 |
| rs976468812 | F175C | Decrease Protein Stability | -1.58 |
| rs398123449 | D196A | Decrease Protein Stability | -1 |
| rs752621035 | R199G | Decrease Protein Stability | -1.64 |
| rs762821794 | G204E | Decrease Protein Stability | -0.94 |
| rs761538803 | R211T | Decrease Protein Stability | -0.74 |
| rs367963796 | L223P | Decrease Protein Stability | -1.81 |
| rs1050332517 | N232H | Decrease Protein Stability | -0.85 |
| rs1375560334 | L234R | Decrease Protein Stability | -1.74 |
| rs1258632902 | H237R | Decrease Protein Stability | -0.16 |
| rs1301523493 | P245L | Decrease Protein Stability | -0.37 |
| rs771103635 | L254S | Decrease Protein Stability | -2.11 |
| rs373979283 | Y266D | Decrease Protein Stability | -1.24 |
| rs398123450 | Y266C | Decrease Protein Stability | -1.41 |
| rs750645495 | E288Q | Decrease Protein Stability | -0.77 |
| rs866283559 | D290N | Decrease Protein Stability | -1.04 |
| rs1200101608 | G293R | Decrease Protein Stability | -0.31 |
| rs1439544901 | W298C | Decrease Protein Stability | -1.35 |
| rs1189912520 | G301D | Decrease Protein Stability | -0.81 |
| rs1252744810 | L305P | Decrease Protein Stability | -1.43 |
| rs773315856 | F344S | Decrease Protein Stability | -1.52 |
| rs751352339 | H350D | Increase Protein Stability | 0.01 |
| rs781244479 | D354A | Decrease Protein Stability | -0.36 |
| rs781244479 | D354G | Increase Protein Stability | -0.86 |
| rs1245724719 | C360Y | Increase Protein Stability | -0.25 |
| rs779364743 | W361R | Decrease Protein Stability | -1.07 |
| rs753450688 | W405R | Decrease Protein Stability | -1.18 |
| rs984639297 | P468H | Decrease Protein Stability | -1.27 |
| rs760333736 | G483D | Decrease Protein Stability | -0.77 |
| rs760333736 | G483V | Decrease Protein Stability | -0.13 |
| rs1447056657 | W489R | Decrease Protein Stability | -1.08 |
| rs754180181 | E491G | Decrease Protein Stability | -1.2 |
| - | V493G | Decrease Protein Stability | -2.58 |
| rs121907985 | P504S | Decrease Protein Stability | -1.7 |
| - | R505W | Decrease Protein Stability | -0.34 |
| rs1229725676 | E511G | Decrease Protein Stability | -1.29 |
| rs778501777 | L513H | Decrease Protein Stability | -2.4 |
| rs778501777 | L513P | Decrease Protein Stability | -1.81 |
| rs1417317737 | W514C | Decrease Protein Stability | -1.66 |
| rs760366178 | R528I | Decrease Protein Stability | -0.39 |
| rs745454291 | L529R | Decrease Protein Stability | -1.77 |
| rs764552042 | R533C | Decrease Protein Stability | -1.05 |
| rs727503960 | C534F | Decrease Protein Stability | 0.04 |
| rs749646826 | R539C | Decrease Protein Stability | -0.79 |
| rs769208361 | R539H | Decrease Protein Stability | -1.15 |
| rs727503961 | C551Y | Decrease Protein Stability | -0.06 |

### 3.4 SNPs effect on 3’UTR Region (miRNA binding sites) in HEXB using PolymiRTS Database

**Table (6):**
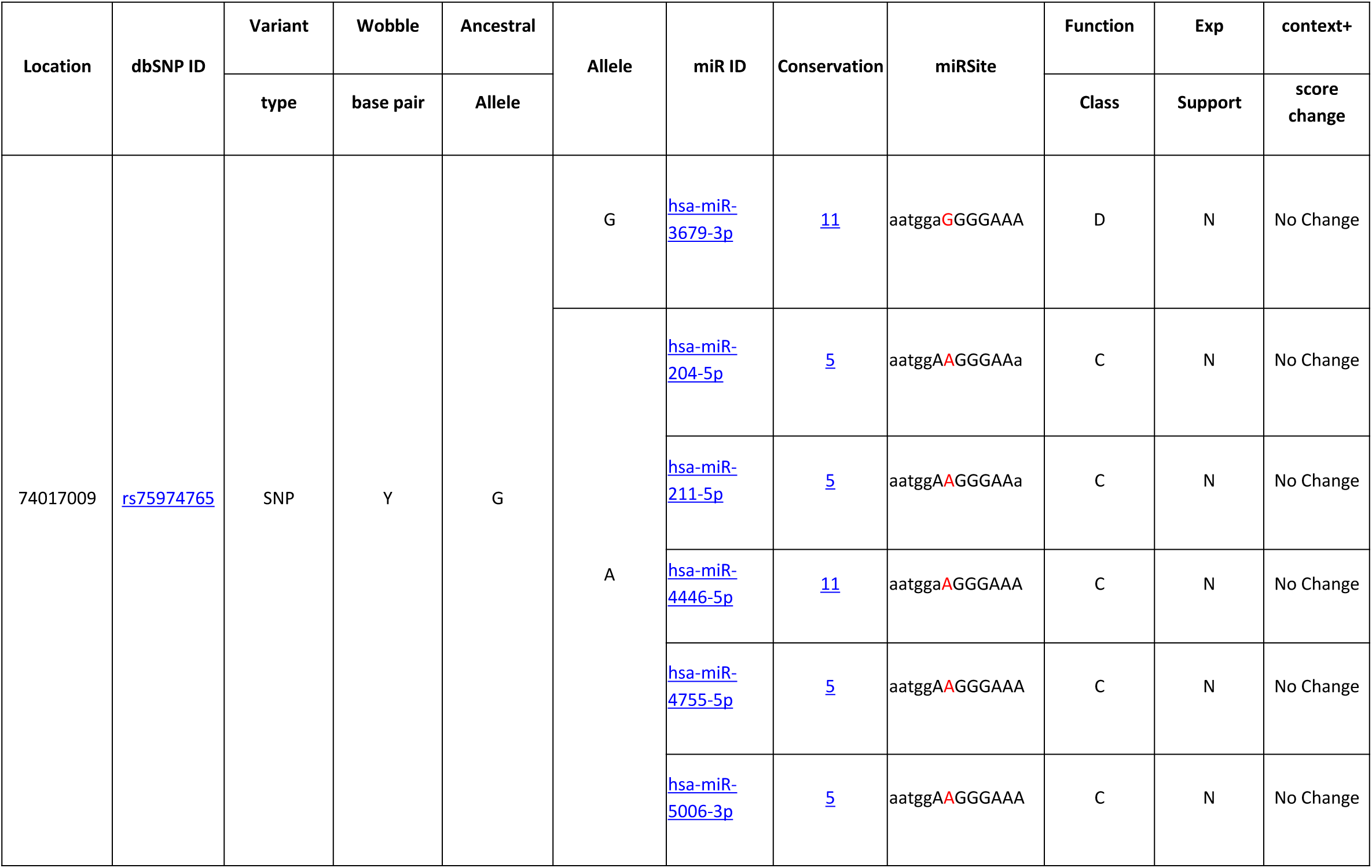
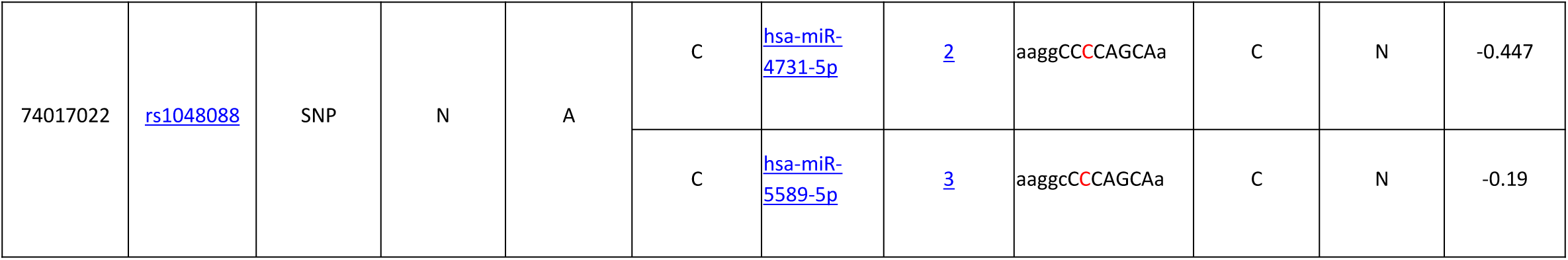
prediction of SNPs sites in HEXB gene at the 3’UTR Region using PolymiRTS

### 3.5 Modeling of amino acid substitution effects on protein structure using

**Figure 2:**
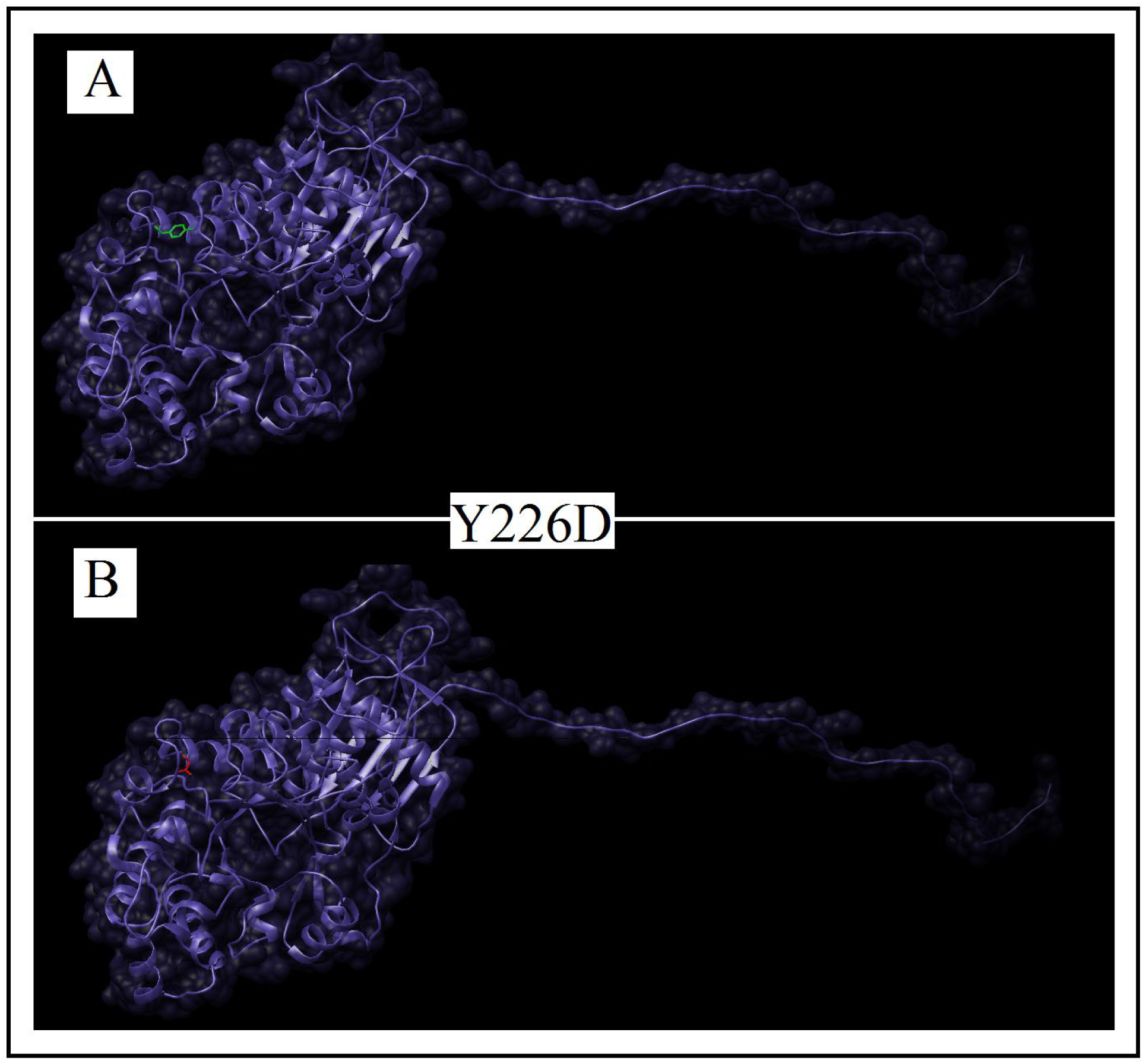
rs373979283 **(Y226D)** Effect of change in the amino acid position **226** from **Threonine** to **Aspartic Acid** on the **HEXB** protein 3D structure using Chimera Software.

**Figure 3:**
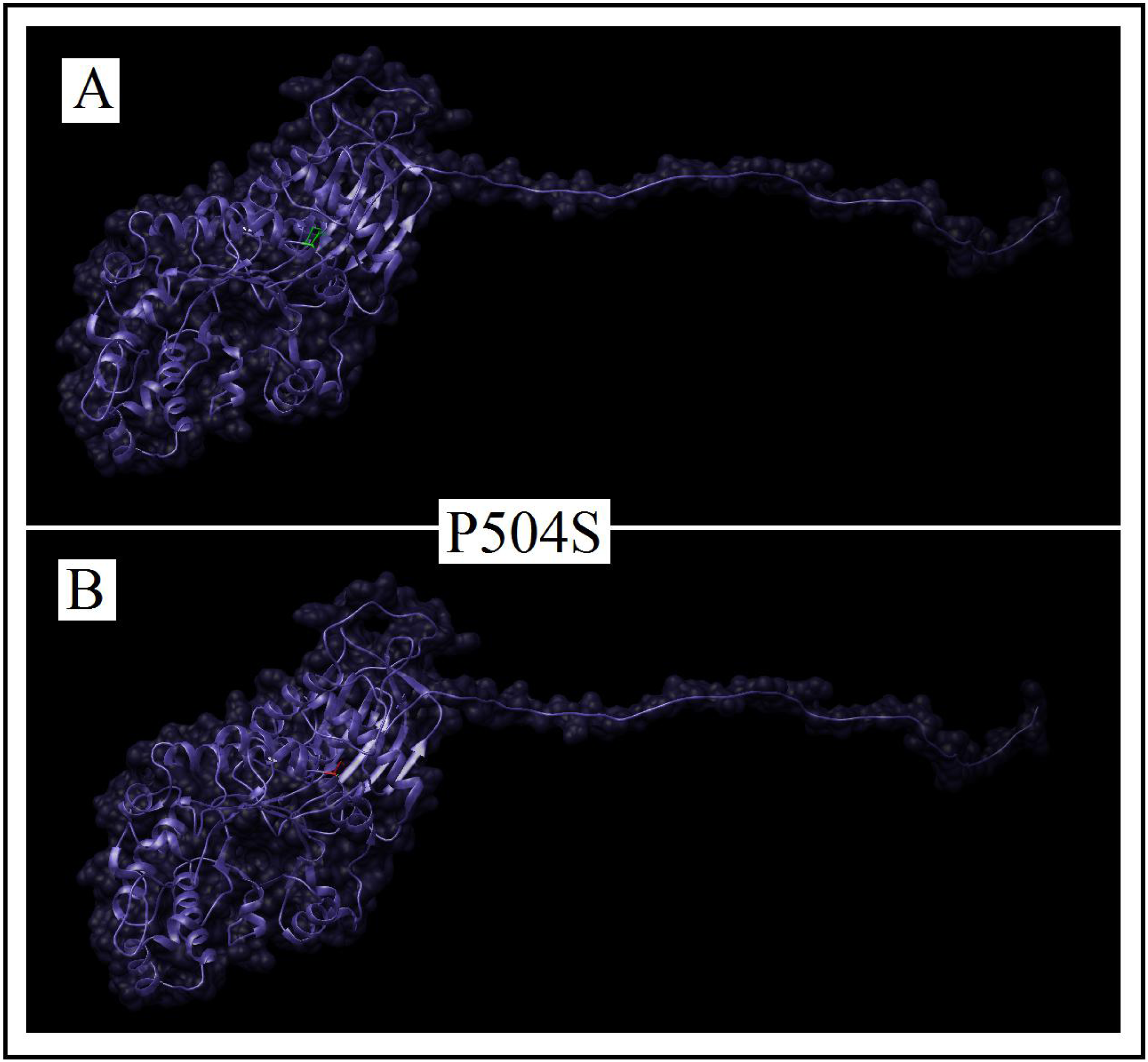
rs121907985 **(P504S)** Effect of change in the amino acid position **504** from **Proline** into **Serine** on the **HEXB** protein 3D structure using Chimera Software.

**Figure 4:**
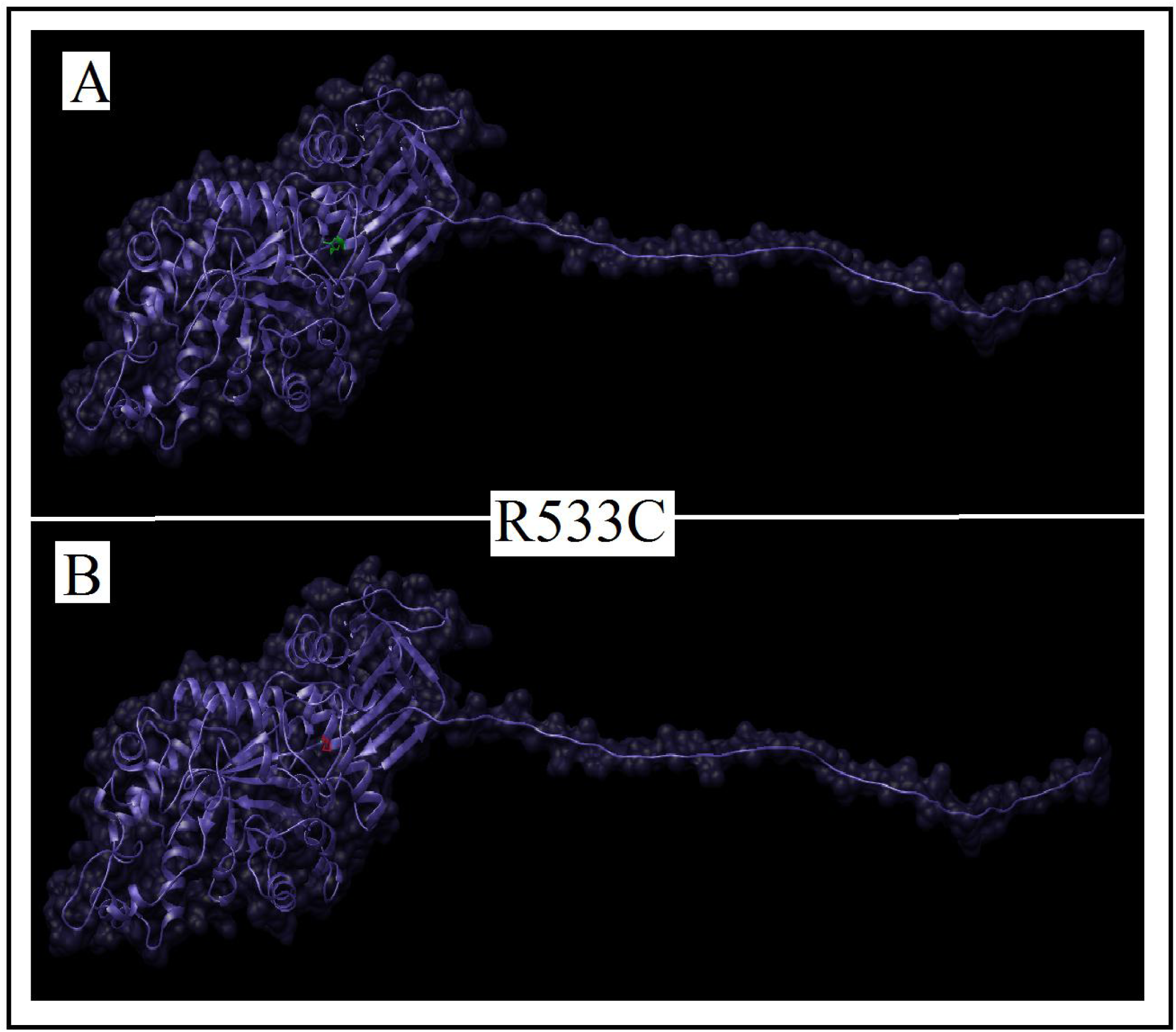
rs764552042 **(R533C)** Effect of change in the amino acid position **533** from **Arginine** into **Cysteine** on the **HEXB** protein 3D structure using Chimera Software.

#### Interactions of *HEXB* gene with other Functional Genes illustrated by GeneMANIA

**Figure 5:**
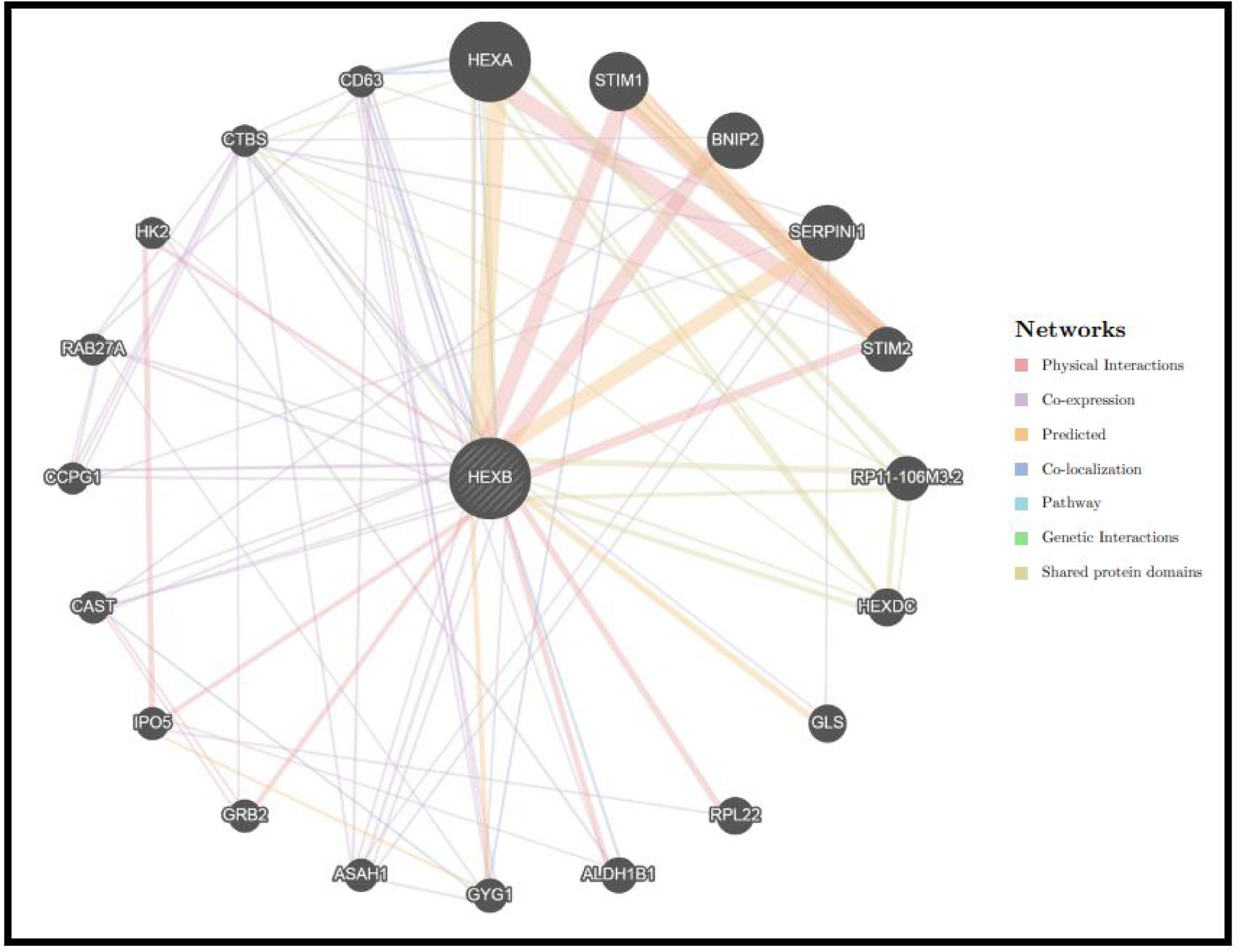
HEXB Gene Interactions and network predicted by GeneMania.

**Table (4):**
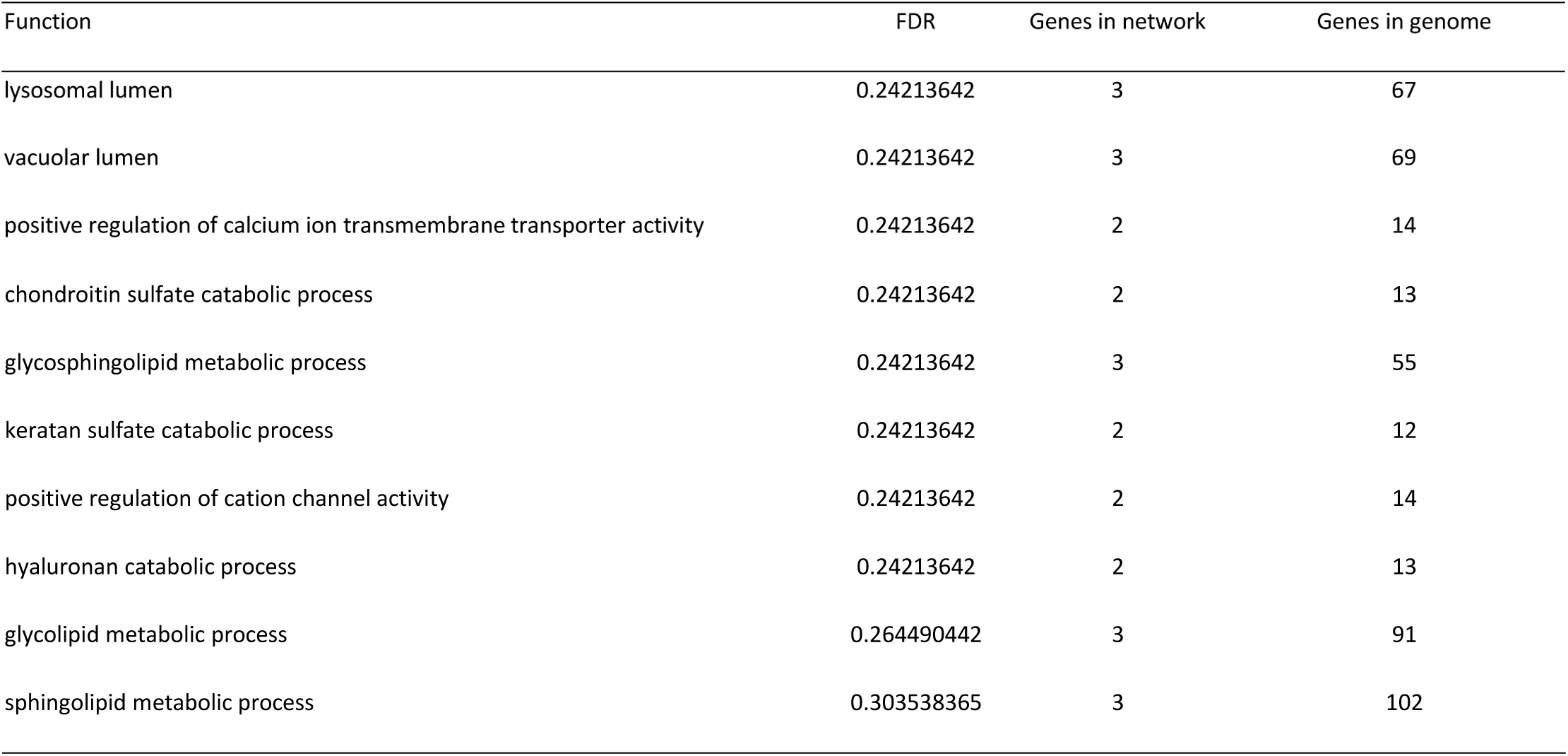
The *HEXB* gene functions and its appearance in network and genome

| Function | FDR | Genes in network | Genes in genome |
| --- | --- | --- | --- |
| lysosomal lumen | 0.24213642 | 3 | 67 |
| vacuolar lumen | 0.24213642 | 3 | 69 |
| positive regulation of calcium ion transmembrane transporter activity | 0.24213642 | 2 | 14 |
| chondroitin sulfate catabolic process | 0.24213642 | 2 | 13 |
| glycosphingolipid metabolic process | 0.24213642 | 3 | 55 |
| keratan sulfate catabolic process | 0.24213642 | 2 | 12 |
| positive regulation of cation channel activity | 0.24213642 | 2 | 14 |
| hyaluronan catabolic process | 0.24213642 | 2 | 13 |
| glycolipid metabolic process | 0.264490442 | 3 | 91 |
| sphingolipid metabolic process | 0.303538365 | 3 | 102 |

**Table (5):** The gene co-expressed, share domain and Interaction with HEXA gene network

| Gene 1 | Gene 2 | Weight | Network group | Network |
| --- | --- | --- | --- | --- |
| *ASAH1* | *HEXB* | 0.024078486 | Co-expression | Ramaswamy-Golub-2001 |
| *CAST* | *HEXB* | 0.013190575 | Co-expression | Ramaswamy-Golub-2001 |
| *RAB27A* | *HEXB* | 0.021869302 | Co-expression | Ramaswamy-Golub-2001 |
| *CD63* | *HEXB* | 0.016662408 | Co-expression | Ramaswamy-Golub-2001 |
| *CD63* | *HEXA* | 0.013731302 | Co-expression | Ramaswamy-Golub-2001 |
| *CD63* | *ASAH1* | 0.016009336 | Co-expression | Ramaswamy-Golub-2001 |
| *CD63* | *RAB27A* | 0.017109819 | Co-expression | Ramaswamy-Golub-2001 |
| *GYG1* | *HEXB* | 0.008671436 | Co-expression | Wang-Maris-2006 |
| *ASAH1* | *HEXB* | 0.01164002 | Co-expression | Wang-Maris-2006 |
| *CAST* | *HEXB* | 0.009363974 | Co-expression | Wang-Maris-2006 |
| *CD63* | *HEXB* | 0.00630279 | Co-expression | Wang-Maris-2006 |
| *CD63* | *GYG1* | 0.011525528 | Co-expression | Wang-Maris-2006 |
| *HEXA* | *HEXB* | 0.021510394 | Co-expression | Mallon-McKay-2013 |
| *CCPG1* | *HEXB* | 0.017047796 | Co-expression | Mallon-McKay-2013 |
| *CTBS* | *CCPG1* | 0.012258756 | Co-expression | Mallon-McKay-2013 |
| *CTBS* | *SERPINI1* | 0.019850705 | Co-expression | Bild-Nevins-2006 B |
| *HK2* | *ALDH1B1* | 0.022080135 | Co-expression | Burington-Shaughnessy-2008 |
| *CTBS* | *GRB2* | 0.007812744 | Co-expression | Burington-Shaughnessy-2008 |
| *CD63* | *SERPINI1* | 0.00854729 | Co-expression | Burington-Shaughnessy-2008 |
| *CD63* | *CTBS* | 0.008849712 | Co-expression | Burington-Shaughnessy-2008 |
| *ASAH1* | *SERPINI1* | 0.012569555 | Co-expression | Dobbin-Giordano-2005 |
| *CCPG1* | *HEXB* | 0.021864697 | Co-expression | Dobbin-Giordano-2005 |
| *RAB27A* | *HEXB* | 0.020799356 | Co-expression | Dobbin-Giordano-2005 |
| *CTBS* | *HEXB* | 0.01395548 | Co-expression | Dobbin-Giordano-2005 |
| *CTBS* | *CCPG1* | 0.017793158 | Co-expression | Dobbin-Giordano-2005 |
| *CTBS* | *RAB27A* | 0.01619344 | Co-expression | Dobbin-Giordano-2005 |
| *CAST* | *HEXB* | 0.01447627 | Co-expression | Bahr-Bowler-2013 |
| *CD63* | *GYG1* | 0.02225664 | Co-expression | Bahr-Bowler-2013 |
| *ASAH1* | *GYG1* | 0.019104764 | Co-expression | Innocenti-Brown-2011 |
| *RAB27A* | *GYG1* | 0.006233362 | Co-expression | Innocenti-Brown-2011 |
| *RAB27A* | *CCPG1* | 0.006827028 | Co-expression | Innocenti-Brown-2011 |
| *IPO5* | *ALDH1B1* | 0.005587121 | Co-expression | Rieger-Chu-2004 |
| *CAST* | *BNIP2* | 0.016995838 | Co-expression | Rieger-Chu-2004 |
| *CD63* | *GYG1* | 0.01405997 | Co-expression | Rieger-Chu-2004 |
| *CTBS* | *HEXB* | 0.011450487 | Co-expression | Noble-Diehl-2008 |
| *CTBS* | *ASAH1* | 0.019146528 | Co-expression | Noble-Diehl-2008 |
| *CCPG1* | *HEXB* | 0.007359452 | Co-expression | Roth-Zlotnik-2006 |
| *RAB27A* | *CCPG1* | 0.018139474 | Co-expression | Roth-Zlotnik-2006 |
| *CTBS* | *HEXB* | 0.005897354 | Co-expression | Roth-Zlotnik-2006 |
| *CTBS* | *CCPG1* | 0.013826602 | Co-expression | Roth-Zlotnik-2006 |
| *CD63* | *HEXB* | 0.003053493 | Co-expression | Roth-Zlotnik-2006 |
| *ASAH1* | *SERPINI1* | 0.008385506 | Co-expression | Perou-Botstein-2000 |
| *CAST* | *HEXB* | 0.015681218 | Co-expression | Perou-Botstein-2000 |
| *CCPG1* | *SERPINI1* | 0.012254277 | Co-expression | Smirnov-Cheung-2009 |
| *CTBS* | *BNIP2* | 0.007756564 | Co-expression | Smirnov-Cheung-2009 |
| *CD63* | *HEXB* | 0.013617347 | Co-expression | Smirnov-Cheung-2009 |
| *IPO5* | *RPL22* | 0.012078757 | Co-expression | Wang-Cheung-2015 |
| *CD63* | *HEXB* | 0.008066914 | Co-expression | Wang-Cheung-2015 |
| *ASAH1* | *HEXB* | 0.023330089 | Co-expression | Chen-Brown-2002 |
| *CTBS* | *STIM2* | 0.011675326 | Co-expression | Perou-Botstein-1999 |
| *GLS* | *HEXB* | 0.005058561 | Co-expression | Wu-Garvey-2007 |
| *GLS* | *SERPINI1* | 0.014305156 | Co-expression | Wu-Garvey-2007 |
| *ALDH1B1* | *HEXB* | 0.006169923 | Co-expression | Wu-Garvey-2007 |
| *HK2* | *HEXB* | 0.005873893 | Co-expression | Wu-Garvey-2007 |
| *CD63* | *HEXA* | 0.010798354 | Co-localization | Schadt-Shoemaker-2004 |
| *CD63* | *ALDH1B1* | 0.00954906 | Co-localization | Schadt-Shoemaker-2004 |
| *HEXA* | *HEXB* | 0.005979959 | Co-localization | Johnson-Shoemaker-2003 |
| *GYG1* | *STIM1* | 0.007238485 | Co-localization | Johnson-Shoemaker-2003 |
| *CAST* | *GYG1* | 0.004685127 | Co-localization | Johnson-Shoemaker-2003 |
| *CTBS* | *HEXB* | 0.008465399 | Co-localization | Johnson-Shoemaker-2003 |
| *CD63* | *HEXA* | 0.008026743 | Co-localization | Johnson-Shoemaker-2003 |
| *BNIP2* | *HEXB* | 0.4508404 | Physical Interactions | Kristensen-Foster-2012 |
| *RPL22* | *HEXB* | 0.1588352 | Physical Interactions | Kristensen-Foster-2012 |
| *ALDH1B1* | *HEXB* | 0.12618154 | Physical Interactions | Kristensen-Foster-2012 |
| *IPO5* | *HEXB* | 0.093262665 | Physical Interactions | Kristensen-Foster-2012 |
| *HK2* | *HEXB* | 0.06835952 | Physical Interactions | Kristensen-Foster-2012 |
| *HK2* | *IPO5* | 0.121480554 | Physical Interactions | Kristensen-Foster-2012 |
| *STIM1* | *HEXB* | 0.6925315 | Physical Interactions | Wan-Emili-2015 |
| *STIM2* | *HEXB* | 0.3523203 | Physical Interactions | Wan-Emili-2015 |
| *STIM2* | *HEXA* | 0.7962252 | Physical Interactions | Wan-Emili-2015 |
| *GRB2* | *HEXB* | 0.14996375 | Physical Interactions | Wan-Emili-2015 |
| *STIM2* | *STIM1* | 0.8585416 | Physical Interactions | BIOGRID-SMALL-SCALE-STUDIES |
| *STIM2* | *STIM1* | 0.2683295 | Physical Interactions | IREF-INTACT |
| *CAST* | *GRB2* | 0.010464651 | Physical Interactions | IREF-INTACT |
| *CAST* | *GRB2* | 0.016936215 | Physical Interactions | Wu-Li-2007 |
| *STIM2* | *STIM1* | 1 | Physical Interactions | IREF-HPRD |
| *HEXA* | *HEXB* | 0.76536685 | Predicted | Wu-Stein-2010 |
| *STIM2* | *STIM1* | 1 | Predicted | Wu-Stein-2010 |
| *SERPINI1* | *HEXB* | 0.8312539 | Predicted | Stuart-Kim-2003 |
| *GLS* | *HEXB* | 0.29914758 | Predicted | Stuart-Kim-2003 |
| *GYG1* | *HEXB* | 0.5665604 | Predicted | I2D-IntAct-Fly2Human |
| *IPO5* | *GYG1* | 0.17434177 | Predicted | I2D-IntAct-Fly2Human |
| *HEXA* | *HEXB* | 0.1421952 | Shared protein domains | INTERPRO |
| *STIM2* | *STIM1* | 0.03625844 | Shared protein domains | INTERPRO |
| *RP11-106M3.2* | *HEXB* | 0.1104058 | Shared protein domains | INTERPRO |
| *RP11-106M3.2* | *HEXA* | 0.1104058 | Shared protein domains | INTERPRO |
| *HEXDC* | *HEXB* | 0.05014918 | Shared protein domains | INTERPRO |
| *HEXDC* | *HEXA* | 0.05014918 | Shared protein domains | INTERPRO |
| *HEXDC* | *RP11-106M3.2* | 0.05154457 | Shared protein domains | INTERPRO |
| *CTBS* | *HEXB* | 0.013065673 | Shared protein domains | INTERPRO |
| *CTBS* | *HEXA* | 0.013065673 | Shared protein domains | INTERPRO |
| *CTBS* | *RP11-106M3.2* | 0.013429452 | Shared protein domains | INTERPRO |
| *CTBS* | *HEXDC* | 0.014957326 | Shared protein domains | INTERPRO |
| *HEXA* | *HEXB* | 0.3798553 | Shared protein domains | PFAM |
| *STIM2* | *STIM1* | 0.05877976 | Shared protein domains | PFAM |
| *RP11-106M3.2* | *HEXB* | 0.3798553 | Shared protein domains | PFAM |
| *RP11-106M3.2* | *HEXA* | 0.3798553 | Shared protein domains | PFAM |
| *HEXDC* | *HEXB* | 0.28301316 | Shared protein domains | PFAM |
| *HEXDC* | *HEXA* | 0.28301316 | Shared protein domains | PFAM |
| *HEXDC* | *RP11-106M3.2* | 0.28301316 | Shared protein domains | PFAM |

## 4. DISSCUTION

42 novel nsSNPs were identified out of 49 most damaging nsSNPs in the *HEXB* gene retrieved by our analysis using different bioinformatics tools which they could have adverse effects on the resulting protein fractionally and structurally.

The variant rs373979283 which entails the substitution Y266D was previously reported in patients with Sandoff disease(41-43), was predicted to be deleterious by all of the softwares used in this study in spite of the fact that Tyrosine is not highly conserved at this position because other homologous sequences were observed having other residues at the same position, but none of them is similar to Aspartate in terms of physiochemical properties, because Aspartate is smaller and less hydrophobic than Tyrosine, which could affect the protein folding process, in addition to that Tyrosine is neutral while Aspartate is negative in charge, therefore, this mutation could still be damaging.

The rs121907985 variant results in converting Proline into Serine which is smaller and less hydrophobic amino acid at the position 504, the difference in size could cause an empty space in the core of the protein, and the internal hydrophobic interactions could be lost because of the difference in the hydrophobicity value. Moreover, Proline’s rigidity could be crucial to induce a special backbone conformation at this position which can be disturbed by this substitution. This mutation is annotated to be pathogenic in ClinVar and ExPASy databases, and was previously reported by Hou *et al.(1998)* in two sisters with Sandoff disease, they found that this mutation significantly increases precursor/mature ratio as a result of increasing the retention of newly synthesized protein in the Endoplasmic reticulum, and also lowers its heat stability(44). The P504S substitution is mostly linked to chronic and less severe forms of the disease (45).

The variant rs764552042 marks the substitution from Arginine to Cysteine at the position 533 (R533C), the mutant residue is neutral in charge, smaller and less hydrophobic than the wild residue which is positive in charge. The difference in charge and size could possibly have some detrimental effects on external interactions and interactions with internal molecules. The same mutation was previously reported in many countries in patients with Sandoff disease (13, 46-48). Zampieri *et al.(2012)* found that the activity of the affected protein is considerably diminished or completely absent compared to the normal protein, and they predicted the mutant residue C533 could form a disulphide bond with C551 which normally forms this bond with C534, and that could result in severe misfolding of the C terminal loop of the protein(49).

All the three mentioned SNPs reside at <u>IPR025705</u> and <u>IPR017853</u> domains, in addition to <u>IPR015883</u> for Y266D, P504S and R533C. These three domains are crucial for normal protein function and they interact with each other, which mean any mutation with deleterious effects within those domains could directly disturb the normal protein functionality.

Two SNPs (<u>rs75974765</u> and <u>rs1048088</u>) were found to have an effect in 3’UTR. rs75974765 SNP is predicted to disrupt binding site of hsa-miR-3679-3p miRNA and creating new binding site at hsa-miR-204-5p, hsa-miR-211-5p, hsa-miR-4446-5p, hsa-miR-4755-5p and hsa-miR-5006-3p miRNAs while <u>rs1048088</u> SNP is predicted to creat new binding site at hsa-miR-4731-5p hsa-miR-5589-5p miRNAs. Both of those SNPs might result in expression of the gene.

The importance of this study relies on the fact that it has subjected all the reported variants within the *HEXB* gene for computational functional and structural analysis which might facilitate further association studies. However, there are some limitations to be considered, like the highly selective protocol which might lead to missing some important SNPs. Computational analysis provides a good insight into which SNPs could drastically affect the protein’s tertiary structure and function, but it’s not completely conclusive, as such, we recommend following this study with wet-lab functional genetic studies and gene knockout on real animal models.

## 5. CONCLUSION

This study revealed 42 novel nsSNP out of 49 damaging SNPs in the *HEXB* gene affecting its function, structure and stability which they possibly lead to Sandhoff disease, by using different bioinformatics tools. Also, 2 SNPs found to have effect on miRNAs binding site affecting expression of HEXB gene. Optimistically, these results will aid in genetic studying and diagnosis of Sandhoff disease improvement.

## 6. ACKNOWLEDGMENT

The authors desire to acknowledge the exciting collaboration of Africa City of Technology – Sudan.

## 7. DATA AVAILABILITY

All relevant data used to support the findings of this study are included within the manuscript and supplementary information files.

